# The transferability of lipid loci across African, Asian and European cohorts

**DOI:** 10.1101/525170

**Authors:** Nikita Telkar, Theresa Reiker, Robin G. Walters, Kuang Lin, Anders Eriksson, Deepti Gurdasani, Arthur Gilly, Lorraine Southam, Emmanouil Tsafantakis, Maria Karaleftheri, Janet Seeley, Anatoli Kamali, Gershim Asiki, Iona Y. Millwood, Michael Holmes, Huaidong Du, Yu Guo, Understanding Society Scientific Group, Meena Kumari, George Dedoussis, Liming Li, Zhengming Chen, Manjinder S. Sandhu, Eleftheria Zeggini, Karoline Kuchenbaecker

## Abstract

The majority of genetic studies for cardiometabolic traits were based on samples with European ancestry. Our aim was to assess whether genetic variants associated with blood lipids, a major risk factor for CVD, are shared across different populations.

We compared genetic associations with lipids between samples from Uganda (N=6,407), China (N=21,295), Japan (N=162,255), the UK (N=9,961) and Greece (N=3,586). Using simulations, we established trans-ethnic colocalization as a method to distinguish shared from population-specific trait loci.

Genetic correlations for HDL, LDL and triglycerides between European ancestry and Asian cohorts were close to 1. A polygenic score based on established LDL-cholesterol-associated loci from European discovery samples had consistent effects on serum levels in samples from the UK, Uganda and Greek population isolates (r=0.23 to 0.28, p<1.9x10^−14^). Overall, ~75% of the major lipid loci from European discovery studies displayed evidence of replication at p<10^−3^, except triglyceride loci in the Ugandan samples of which only 10% replicated. Specific replicating loci were identified using trans-ethnic colocalization. Ten of the fourteen lipid loci that did not replicate in the Ugandan population had pleiotropic associations with BMI in European ancestry samples while none of the replicating loci did. While lipid associations were highly consistent across European and Asian populations, there was a lack of replication particularly for established triglyceride loci in the Ugandan population. These loci might affect lipids by modifying food intake or metabolism in an environment offering diets rich in certain nutrients. This suggests that gene-environment interactions could play an important role for the transferability of complex trait loci.

## Introduction

Cardiovascular disease (CVD) is one of the leading causes of death worldwide^1^. As the predictive ability of common variants for CVD and cardiometabolic traits improves, risk prediction in clinical settings finds increasing consideration^2,3^. The foundations for this were provided by genome “white” association studies: the majority of samples included in these studies were British or US-Americans with European ancestry^4,5^ which does not accurately representation the ethnically and ancestrally diverse populations of these nations. Moreover, three quarters of CVD deaths occur in low- and middle-income countries with incidences further rising^6^. Consequently, it is important to determine whether cardiometabolic trait loci are transferable to other populations. We focussed on blood lipids, a major cardiovascular risk factor.

Previous research assessed the effects of different allele frequencies and linkage disequilibrium (LD) on genetic associations across ancestry groups^7^. Here we ask the fundamental question whether causal variants for lipid biomarkers are shared across populations. Heterogeneity in effects of variants could result from epistasis or gene-environment interactions. However, differences in LD structure between populations make it difficult to compare associations between ancestry groups because the observable effect of a variant depends on its correlation with the causal variant(s)^7^. Differences in frequency also impact on the power to detect associations in other ancestry groups.

We employed several strategies which account for these effects to quantify the extent to which genetic variants affecting lipid biomarkers are shared between individuals from Europe/North America, Asia, and Africa. We assessed the transferability of individual signals and compared association patterns across the genome using data from the African Partnership for Chronic Disease Research – Uganda (APCDR-Uganda, N=6,407)^8^, China Kadoorie Biobank (CKB, N=21,295)^9^, the Hellenic Isolated Cohorts (HELIC-MANOLIS, N=1,641 and HELIC-Pomak, N=1,945)^10,11^, and the UK Household Longitudinal Study (UKHLS, N=9,961)^12^. We also used summary statistics from Biobank Japan (BBJ, N=162,255)^13^ and the Global Lipid Genetics Consortium (European ancestry, GLGC2013 N=188,577, GLGC2017 N=237,050)^14,15^.

## Results

We assessed replication rates across established lipid-associated variants in different populations. We distinguished major lipid loci, i.e. those with p<10^−100^ in the largest European ancestry GWAS. In this context, replication was operationalised as at least one variant from the credible set associated at p<10^−3^ in the target study. As a benchmark, we also assessed replication in two European ancestry studies. We found evidence of replication for 76.5% of major HDL loci in these two studies (Table 1). For the non-European groups replication rates ranged from 70.6 to 82.4%. Similar replication rates were observed for LDL loci (61.5-76.9%). For major triglycerides (TG) loci, replication rates ranged from 78.9 to 94.7%, except in APCDR-Uganda. Only 10.5% of these loci showed evidence of replication in that sample. Replication rates for known loci with p≥10^−100^ in the discovery set were generally moderate to low. However, Biobank Japan, the largest study, had markedly higher replication rates for these loci than the other studies.

**Table 1.**
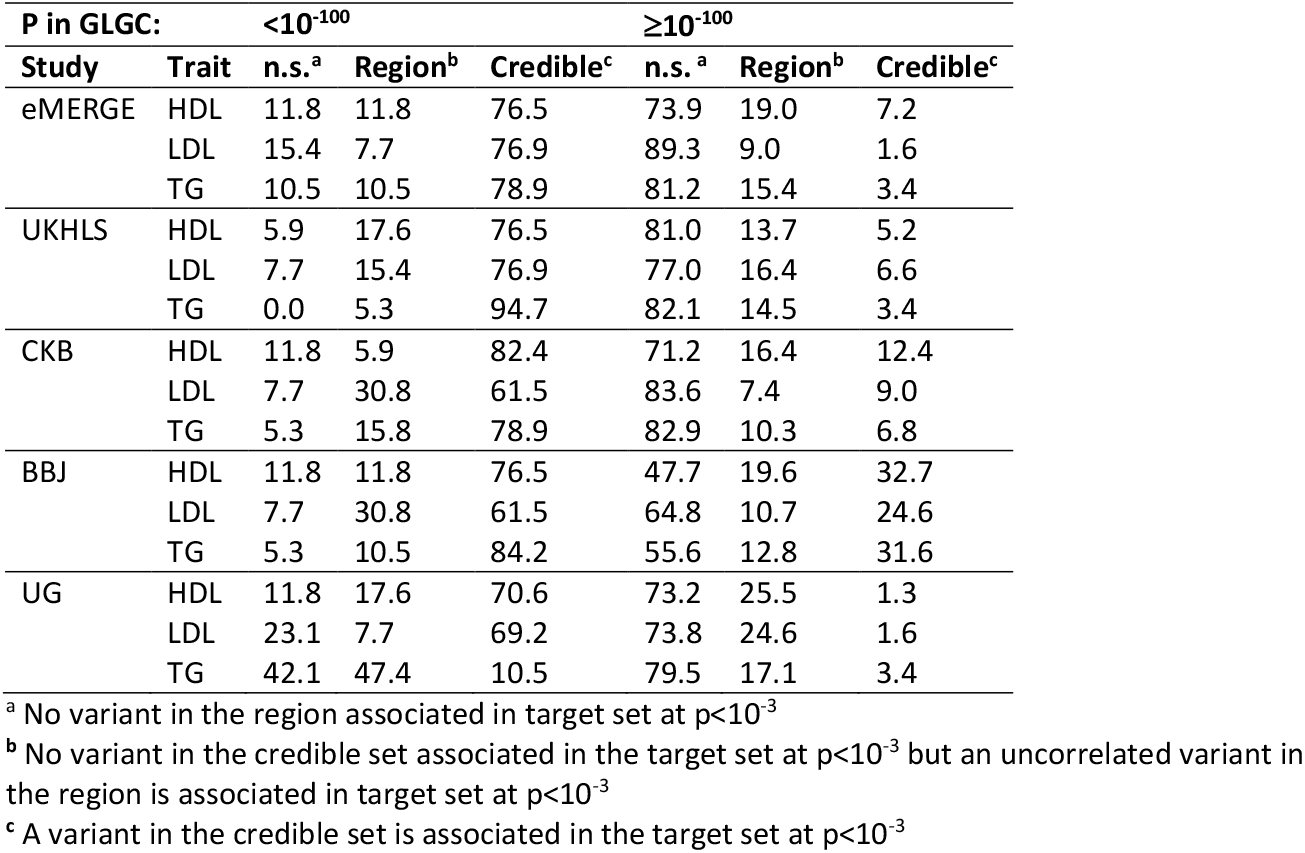
**Percentage of established lipid-associated loci with evidence of replication** in each target study. Results shown separately by strength of association (whether p<10^−100^) in the discovery study (GLGC). Only one SNP was kept for each locus with multiple associated variants in close proximity. Regions were defined as 25Kb either side of the lead variant. The credible set contains the reported lead variant and variants in LD (r2>0.6) with it.

| P in GLGC: |  | <10<sup>-100</sup> |  |  | ≥10<sup>-100</sup> |  |  |
| --- | --- | --- | --- | --- | --- | --- | --- |
| Study | Trait | n.s.<sup>a</sup> | Region<sup>b</sup> | Credible<sup>c</sup> | n.s.<sup>a</sup> | Region<sup>b</sup> | Credible<sup>c</sup> |
| eMERGE | HDL | 11.8 | 11.8 | 76.5 | 73.9 | 19.0 | 7.2 |
|  | LDL | 15.4 | 7.7 | 76.9 | 89.3 | 9.0 | 1.6 |
|  | TG | 10.5 | 10.5 | 78.9 | 81.2 | 15.4 | 3.4 |
| UKHLS | HDL | 5.9 | 17.6 | 76.5 | 81.0 | 13.7 | 5.2 |
|  | LDL | 7.7 | 15.4 | 76.9 | 77.0 | 16.4 | 6.6 |
|  | TG | 0.0 | 5.3 | 94.7 | 82.1 | 14.5 | 3.4 |
| CKB | HDL | 11.8 | 5.9 | 82.4 | 71.2 | 16.4 | 12.4 |
|  | LDL | 7.7 | 30.8 | 61.5 | 83.6 | 7.4 | 9.0 |
|  | TG | 5.3 | 15.8 | 78.9 | 82.9 | 10.3 | 6.8 |
| BBJ | HDL | 11.8 | 11.8 | 76.5 | 47.7 | 19.6 | 32.7 |
|  | LDL | 7.7 | 30.8 | 61.5 | 64.8 | 10.7 | 24.6 |
|  | TG | 5.3 | 10.5 | 84.2 | 55.6 | 12.8 | 31.6 |
| UG | HDL | 11.8 | 17.6 | 70.6 | 73.2 | 25.5 | 1.3 |
|  | LDL | 23.1 | 7.7 | 69.2 | 73.8 | 24.6 | 1.6 |
|  | TG | 42.1 | 47.4 | 10.5 | 79.5 | 17.1 | 3.4 |

<sup>a</sup> No variant in the region associated in target set at $p < 10^{-3}$
<sup>b</sup> No variant in the credible set associated in the target set at $p < 10^{-3}$ but an uncorrelated variant in the region is associated in target set at $p < 10^{-3}$
<sup>c</sup> A variant in the credible set is associated in the target set at $p < 10^{-3}$

Trans-ethnic genetic correlations were estimated between the three largest studies, China Kadoorie Biobank, Biobank Japan and GLGC2013. Correlations were high for each biomarker and were not significantly different from 1 (Figure 1, Supplementary Table 1). We also compared associations across biomarkers. This consistently showed negative genetic correlations between TG associations and HDL associations, with estimates ranging from r_gen_=−0.48 to r_gen_=−0.86.

**Figure 1.**
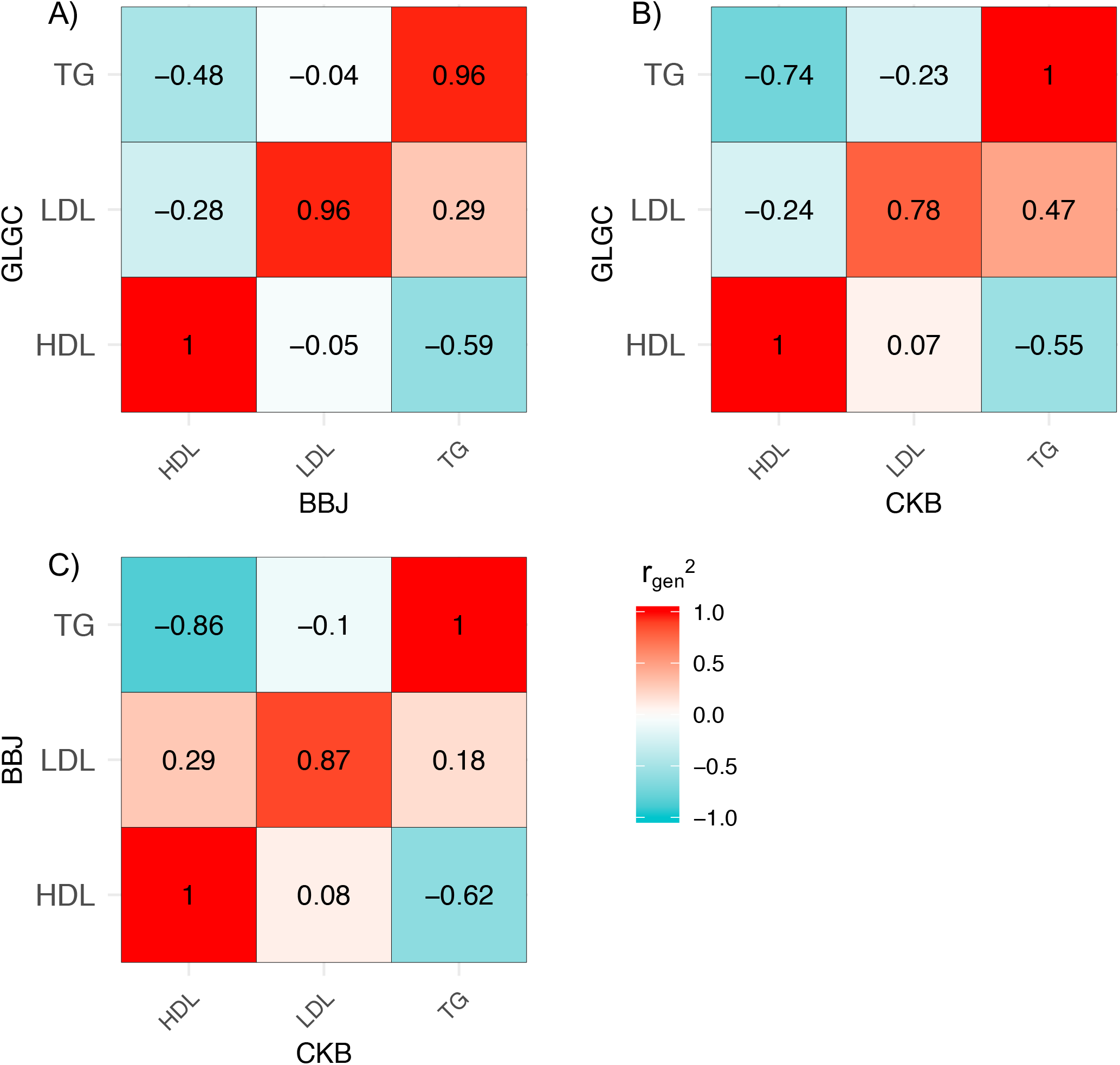
**Trans-ethnic genetic correlations** for associations with high-density lipoprotein (HDL), low-density lipoprotein (LDL) cholesterol and triglycerides (TG). a) shows the comparison of GLGC2013 (European) and Biobank Japan, b) GLGC2013 and China Kadoorie Biobank and c) Biobank Japan and China Kadoorie Biobank.

In order to assess patterns of sharing of risk alleles for the smaller studies, we constructed polygenic scores based on the established lipid loci from discovery samples with European ancestry and estimated the score associations with levels of HDL, LDL and TG in HELIC, APCDR-Uganda and also UKHLS as a benchmark (Figure 2). All genetic scores were significantly associated with their respective target lipid in the three European samples with largely consistent correlation coefficients and mutually overlapping 95% confidence intervals (CIs) (Table 2). For HDL, LDL and TG, the estimated correlation coefficients had a range of 0.27-0.28, 0.23-0.28 and 0.20-0.24, respectively. In APCDR-Uganda, the strongest association was observed for LDL (r=0.28, SE=0.01, p=1.9x10^−107^). The HDL association was attenuated compared to the European samples (r=0.12, SE=0.01, p=6.1x10-^22^). The effect of the TG score was markedly weaker (r=0.06, SE=0.01, p=4.5x10^−7^). We also assessed associations between a given score and levels of each of the other biomarkers (Supplementary Table 2). In line with the trans-ethnic genetic correlation results, we observed inverse associations between the HDL score and TG levels and vice versa in all studies, except APCDR-Uganda.

**Figure 2.**
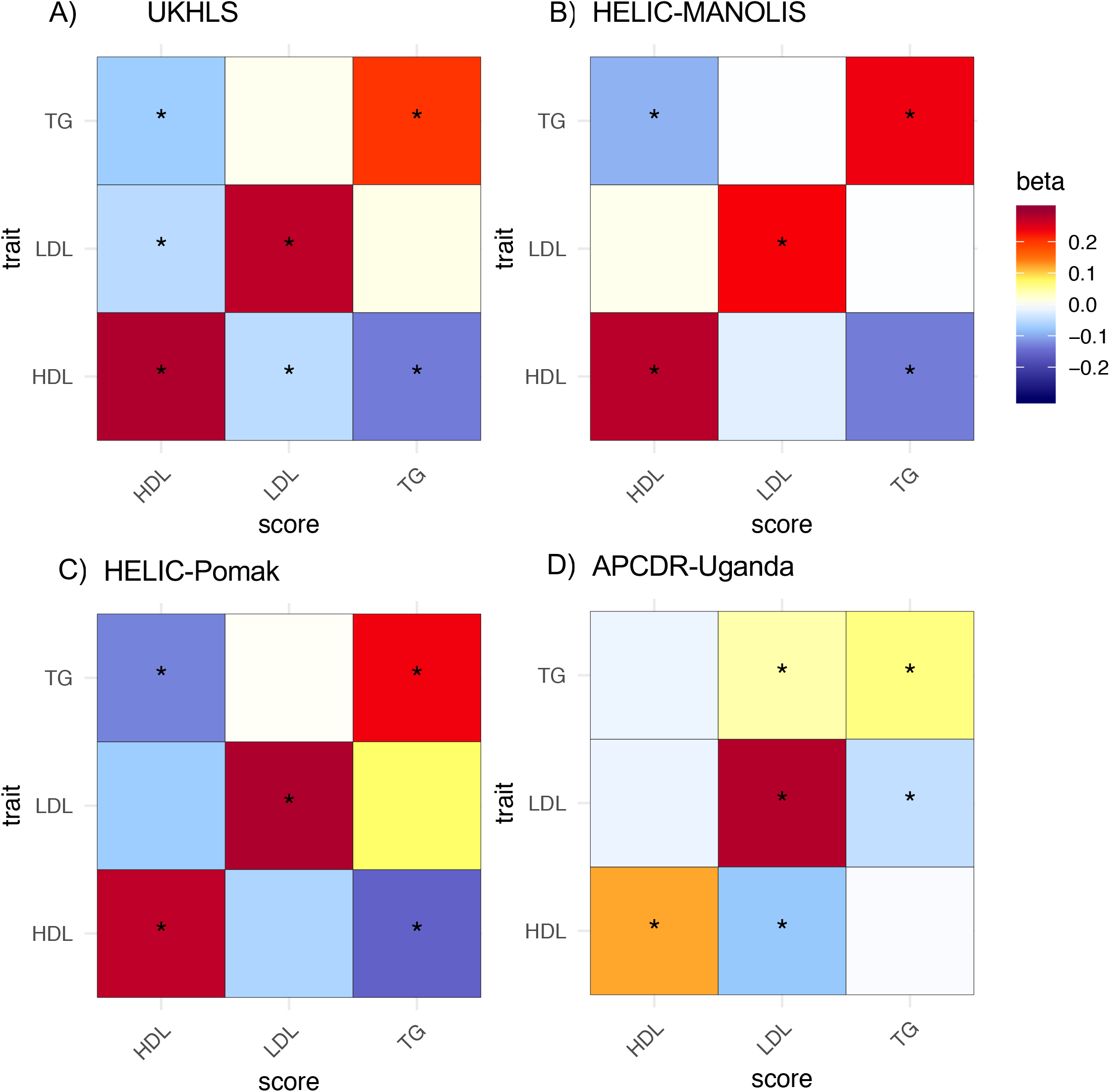
**Associations of polygenic scores** based on established lipid-associated loci with levels of high-density lipoprotein (HDL), low-density lipoprotein (LDL) cholesterol and triglycerides (TG) in a) UKHLS, b) HELIC-MANOLIS, c) HELIC-Pomak, d) APCDR-Uganda. Estimates are given as correlation coefficients. Stars indicate statistically significant associations (p<0.0056).

**Table 2:** **Associations of polygenic scores** based on established lipid-associated loci and respective biomarkers levels in UKHLS, HELIC-MANOLIS, -Pomak, and APCDR-Uganda using a linear mixed model analysis.

| Trait | N | Correlation (SE<sup>a</sup>) | P-value |
| --- | --- | --- | --- |
| **UKHLS** |  |  |  |
| HDL | 9706 | 0.284 (0.010) | 8.34x10<sup>-165</sup> |
| LDL | 9767 | 0.273 (0.010) | 8.38x10<sup>-155</sup> |
| Triglycerides | 9635 | 0.203 (0.010) | 2.62x10<sup>-86</sup> |
| **HELIC-MANOLIS** |  |  |  |
| HDL | 1186 | 0.276 (0.029) | 8.65x10<sup>-20</sup> |
| LDL | 1186 | 0.230 (0.029) | 1.89x10<sup>-14</sup> |
| Triglycerides | 1176 | 0.237 (0.030) | 3.01x10<sup>-14</sup> |
| **HELIC-Pomak** |  |  |  |
| HDL | 1078 | 0.272 (0.030) | 9.67x10<sup>-18</sup> |
| LDL | 1075 | 0.285 (0.030) | 1.35x10<sup>-18</sup> |
| Triglycerides | 1066 | 0.235 (0.030) | 1.68x10<sup>-13</sup> |
| **APCDR-Uganda** |  |  |  |
| HDL | 6407 | 0.121 (0.012) | 6.06x10<sup>-22</sup> |
| LDL | 6407 | 0.280 (0.012) | 1.91x10<sup>-107</sup> |
| Triglycerides | 6407 | 0.063 (0.013) | 4.46x10<sup>-7</sup> |

<sup>a</sup> SE=standard error

Differences in LD structure, MAF and sample size make it difficult to assess replication for individual loci. Therefore, we propose a new strategy to assess evidence for shared causal variants between two populations: trans-ethnic colocalization. For this we re-purposed a method that was originally developed for colocalization of GWAS and eQTL results: Joint Likelihood Mapping (JLIM)^16^. In order to assess its performance for GWAS results from samples with different ancestry, we carried out a simulation study. UK Biobank (UKB) was used as a European ancestry reference and compared to CKB and APCDR-Uganda. Phenotypes were simulated. In the simulations of distinct causal variants in the non-European and the reference group, the frequencies of false negatives were close to 0.05 as expected (Table 3), with an almost uniform distribution of p-values (Supplementary Figure 1). The power to detect shared associations was good for both populations: 73.1% for APCDR-Uganda and 93.1% for CKB. To investigate whether the lower power for APCDR-Uganda could be due to its smaller sample size, we reran the analyses for CKB using a random subset of samples matching the sample size of APCDR-Uganda. The results were similar to the ones for the full CKB set, suggesting that the power of this trans-ethnic colocalization method decreases somewhat with greater genetic distance between the populations that are compared.

**Table 3:**
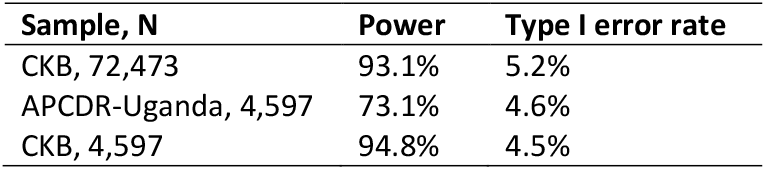
**10,000 simulation runs to assess the performance of trans-ethnic colocalization.** Phenotypes were simulated for CKB and APCDR-Uganda and trans-ethnic colocalization was run to compare each to a reference set of 50,000 samples with British ancestry from UK Biobank.

We applied trans-ethnic colocalization for established lipid loci to each study with UKHLS as the reference. There was evidence for significant (p_jlim_<0.05) colocalization with at least one of the target studies for about half of the major lipid loci (Supplementary Table 3). For several major TG loci, such as *GCKR* at 2p23.3 or *LPL* at 8p21.3, strong evidence of replication in the Asian studies was observed while there was no evidence of association in APCDR-Uganda (Figure 3b,c).

**Figure 3.**
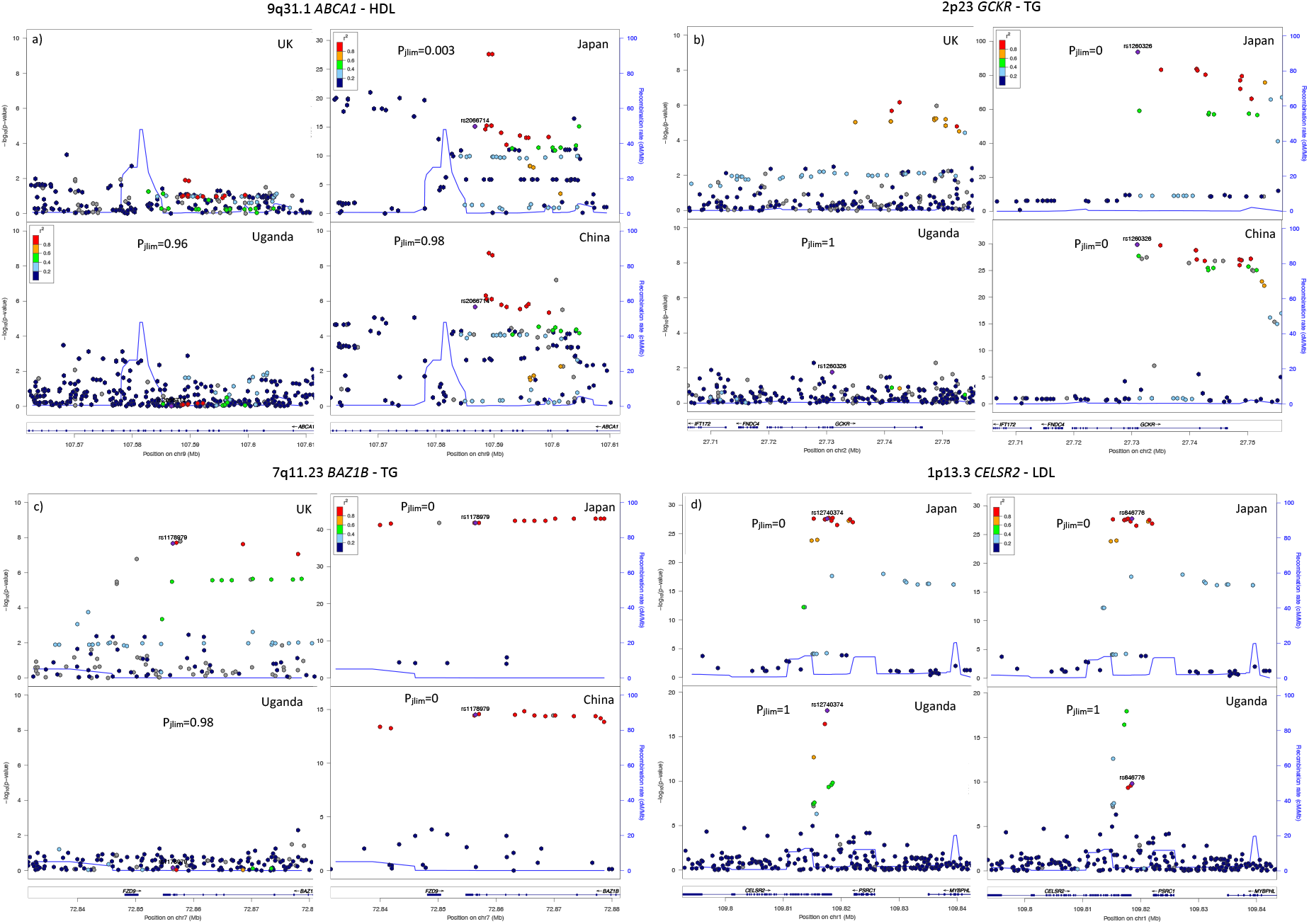
**Regional association plots** for a selection of established lipid-associated loci for UKHLS, Biobank Japan, APCDR-Uganda, and China Kadoorie Biobank and p-value p_jlim_ for the trans-ethnic colocalization with UKHLS.

We compared major lipid loci showing evidence of replication in APCDR-Uganda with those not displaying any suggestion of replication. The proximal genes of replicating loci were enriched for lipid pathways including lipoprotein metabolism, lipid digestion mobilisation and transport, chylomicron-mediated lipid transport and metabolism of lipids and lipoproteins. The proximal genes of the non-replicating loci were enriched for several other pathways in addition to lipid metabolism, including SHP2 signalling, ABV3 integrin pathway, cytokine signalling in immune system, cytokine-cytokine receptor interaction and transmembrane transport of small molecules (Supplementary Figures 2 and 3). We also assessed the associations of these loci with BMI in samples with European ancestry using publicly available summary statistics from the GIANT consortium^17^ (N≥484,680) (Table 4). Ten of the fourteen non-replicating lipid loci had pleiotropic associations with BMI at a Bonferroni-adjusted threshold of p<0.0024. None of the seven replicating lipid loci were associated with BMI. We also assessed four additional loci that were not significant in the trans-ethnic colocalization but displayed small regional p-values in APCDR-Uganda. Out of these only APOE was significantly associated with BMI (p=4x10^−21^).

**Table 4.** **Association of established lipid-associated loci with body mass index** by locus replication status in APCDR-Uganda. Association results are based on N≥484,680 samples from the meta-analysis between GIANT and UK Biobank

| Replication | Trait | rs-id | Chr | Position | Annotation | beta | SE | P-value |
| --- | --- | --- | --- | --- | --- | --- | --- | --- |
| no | HDL | rs11755393 | 6 | 34824636 | *UHRF1BP1* | -0.025 | 0.002 | **9.8x10<sup>-48</sup>** |
| no | HDL | rs1178979 | 7 | 72856430 | *BAZ1B* | -0.010 | 0.002 | **3.1x10<sup>-6</sup>** |
| no | HDL | rs4731702 | 7 | 130433384 | *KLF14* | 0.008 | 0.002 | **3x10<sup>-7</sup>** |
| no | HDL | rs2954033 | 8 | 126493746 | *NSMCE2* | -0.010 | 0.002 | **6.4x10<sup>-8</sup>** |
| no | LDL | rs4245791 | 2 | 44074431 | *ABCG8* | 0.002 | 0.002 | 0.22 |
| no | LDL | rs3846662 | 5 | 74651084 | *HMGCR* | 0.020 | 0.002 | **1.9x10<sup>-35</sup>** |
| no | LDL | rs2737229 | 8 | 116648565 | *TRPS1* | 0.014 | 0.002 | **1.9x10<sup>-15</sup>** |
| no | LDL | rs635634 | 9 | 136155000 | *IL6R* | 0.005 | 0.002 | 0.03 |
| no | LDL | rs2000999 | 16 | 72108093 | *HPR* | 0.011 | 0.002 | **8.6x10<sup>-8</sup>** |
| no | TG | rs1260326 | 2 | 27730940 | *GCKR* | -0.011 | 0.002 | **1.2x10<sup>-10</sup>** |
| no | TG | rs2943641 | 2 | 227093745 | *IRS1* | 0.006 | 0.002 | **5.8x10<sup>-4</sup>** |
| no | TG | rs6905288 | 6 | 43758873 | *VEGFA* | -0.010 | 0.002 | **1.9x10<sup>-9</sup>** |
| no | TG | rs11820589 | 11 | 116633862 | *APOA5* | -0.003 | 0.003 | 0.41 |
| no | TG | rs58542926 | 19 | 19379549 | *TM6SF2* | -0.003 | 0.003 | 0.33 |
| yes | HDL | rs643531 | 9 | 15296034 | *TTC39B* | 0.000 | 0.002 | 0.92 |
| yes | HDL | rs1800588 | 15 | 58723675 | *LIPC* | -0.002 | 0.002 | 0.25 |
| yes | HDL | rs3764261 | 16 | 56993324 | *CETP* | -0.002 | 0.002 | 0.39 |
| yes | HDL | rs16942887 | 16 | 67928042 | *PSKH1* | -0.005 | 0.003 | 0.06 |
| yes | LDL | rs12740374 | 1 | 109817590 | *CELSR2* | 0.003 | 0.002 | 0.18 |
| yes | LDL | rs1367117 | 2 | 21263900 | *APOB* | -0.002 | 0.002 | 0.19 |
| yes | LDL | rs6511720 | 19 | 11202306 | *LDLR* | 0.006 | 0.003 | 0.03 |
| suggestive | HDL | rs4841132 | 8 | 9183596 | *PPP1R3B* | 0.000 | 0.003 | 0.88 |
| suggestive | HDL | rs328 | 8 | 19819724 | *LPL* | 0.005 | 0.003 | 0.10 |
| suggestive | HDL | rs769449 | 19 | 45410002 | *APOE* | -0.026 | 0.003 | **4.4x10<sup>-21</sup>** |
| suggestive | HDL | rs386000 | 19 | 54792761 | *LILRB2* | 0.001 | 0.002 | 0.60 |

## Discussion

Recent efforts to increase global diversity in genetics studies have been vital, enabling this comprehensive cross-population comparison of genetic associations with blood lipids. We provide evidence for extensive sharing of genetic variants that affect levels of HDL- and LDL-cholesterol and triglycerides between individuals with European ancestry and samples from China, Japan and Greek population isolates. There was evidence of replication for about three quarters of major HDL and LDL loci (and triglyceride loci except in APCDR-Uganda). This was highly consistent across all studies. Estimates of trans-ethnic genetic correlations between European, Chinese and Japanese samples were close to 1. Associations of polygenic scores for LDL were not attenuated in the Ugandan population compared to the UK samples. All PRS associations in the two Greek isolated populations were also highly consistent with those in the UK samples.

Previous studies that compared the direction of effect of established loci or assessed associations of polygenic scores reported differing degrees of consistency^18–29^. However, most of them were conducted in American samples with diverse ancestry, had smaller sample sizes and applied a single-variant look-up or PRS for a limited number of genetic variants. The high degree of consistency for cholesterol biomarkers we observed also contrasts with previously reported trans-ethnic genetic correlations for other traits, such as major depression, rheumatoid arthritis, or type 2 diabetes, which were substantially different from 1^30,31^. In a recent application using data from individuals with European and Asian ancestry from the UK and USA, the average genetic correlation across multiple traits was 0.55 (SE = 0.14) for GERA and 0.54 (SE=0.18) for UK Biobank^32^.

Differences in LD structure, MAF and sample size make it difficult to assess replication of individual loci. We therefore propose a new approach: trans-ethnic colocalization. Simulations showed consistent control of type I error rates as well as good power to detect associations. Colocalization identified shared causal variants even at loci where none of the individual variants were associated at stringent p-value thresholds. However, for many of the major lipid loci, more than one independent association signal was identified in discovery GWAS^15^. When these are located in close proximity to each other, they can interfere with the trans-ethnic colocalization analysis because JLIM assumes a single causal variant (Figure 3d). Therefore, future work should extend this approach to accommodate loci harbouring multiple causal variants. Using trans-ethnic colocalization, we showed that many established loci for triglycerides did not affect levels of this biomarker in Ugandan samples. This included loci associated at genome-wide significance in all the other studies, such as *GCKR* at 2p23.3 or *LPL* at 8p21.3. The polygenic score for triglycerides had a weak effect on measured levels in APCDR-Uganda. This is unlikely to be an artefact of unreliable measurement: triglyceride levels had a heritable component in this sample (SNP heritability of 0.25, SE=0.05^8^) and there were some genome-wide significant associations (Supplementary Figure 6e). It is also unlikely that this can be explained purely by differences in LD and MAF because they would affect the analyses of the other two biomarkers as well. Instead these discrepancies could be caused by gene-environment interactions. Most of the lipid loci that did not replicate in the Ugandans had pleiotropic associations with BMI in European ancestry samples while none of the replicating loci were linked to BMI. It is possible that the non-replicating variants affect the amount of food intake with downstream consequences for lipid levels. This might require an environment offering diets that are rich in certain nutrients. While the replicating genes were almost exclusively linked to pathways of lipid metabolism, the non-replicating genes were involved in a diverse pathways which is in line with hypothesis. An alternative explanation could be that the non-replicating loci are involved in metabolising nutrients given a particular diet that is not common in Uganda with downstream consequences for weight.

Overall, this could suggest an important role of environmental factors in modifying which genetic variants affect lipid levels. Studying the causes for discordant loci between groups has promise to further elucidate the biological mechanisms of lipid regulation and other complex traits. Applying genetic risk prediction within clinical settings is receiving increasing attention. Our findings demonstrate that the transferability of genetic associations across different ancestry groups and environmental settings should be assessed comprehensively for medically relevant traits. This is important in order to maximise the potential of precision medicine to yield health benefits that are widely shared within and across populations. Ongoing programs in underrepresented countries^33^, such as the Human Hereditary and Health in Africa Initiative^34^, and programs focussing on underrepresented groups, such as PAGE^35^, All of Us^36^, or East London Genes and Health^37^, could provide the basis for this.

## Methods

### Data resources

We included data from the Global Lipid Genetics Consortium (European ancestry samples only, GLGC), The UK Household Longitudinal Study (UKHLS), two isolated populations from the Greece Hellenic Isolated Cohorts (HELIC), a rural West Ugandan population from the African Partnership for Chronic Disease Research (APCDR-Uganda) study, China Kadoorie Biobank (CBK), and Biobank Japan (BBJ). In addition, we used data from European ancestry samples from the eMERGE network to confirm replication rates of known loci. Raw genotype and phenotype data were available for UKHLS, APCDR-Uganda, CKB, HELIC-MANOLIS, HELIC-Pomak and eMERGE. Our analyses were based on summary statistics for BBJ and GLGC. Study details are provided in Supplementary Table 3.

Each study underwent standard quality control. Details of the genome-wide association analyses with lipid traits have been previously described for GLGC^14^, BBJ^13^, HELIC^10^, and UKHLS^12^. The association analysis for APCDR-Uganda was carried out within a mixed model framework using GEMMA^38^. Rank-based inverse normal transformation was applied to the lipid biomarkers after adjusting for age and gender. In China Kadoorie Biobank, lipid levels were regressed against eight principle components, region, age, age^2^, sex, and - for LDL and TG - fasting time^2^. LDL levels were derived using the Friedewald formula. After rank-based inverse normal transformation, the residuals were used as the outcomes in the genetic association analyses using linear regression. In eMERGE, biomarkers were adjusted for age, gender, kidney disease, statin use, type 2 diabetes status and disorders relating to growth hormones. Associations were carried out within a mixed model framework using BOLT-LMM^39^. Manhattan plots for eMerge, UKHLS, BBJ, CKB, and APCDR-Uganda are shown in Supplementary Figures 4-6.

### Established lipid loci

A list of established lipid-associated loci was extracted from the latest Global Lipid Genetics Consortium (GLGC2017) publication^15^ reporting 444 independent variants in 250 loci associated at genome-wide significance with HDL, LDL, and triglyceride levels. We excluded three LDL variants where the association was not primarily driven by the samples with European ancestry. We assessed evidence of replication of the loci, applied trans-ethnic colocalization and used them to construct polygenic scores.

### Replication of established lipid loci

We assessed evidence of replication across these established lipid variants. For loci harbouring multiple signals, we only kept the most strongly associated variant. This left 170 HDL, 135 LDL and 136 TG variants. We distinguished major loci, i.e. those with p<10^−100^ in GLGC2017. For each lead SNP we identified all variants in LD (r^2^>0.6) based on the European ancestry 1000 Genomes data. We assessed whether the lead or any of the correlated variants, henceforth called credible set, displayed evidence of association in the target study. We used a p-value threshold of p<10^−3^. If this was not the case, we tested whether there was any other variant with evidence of association within a 50Kb window. While this p-value threshold might not be appropriate to provide conclusive evidence of replication for individual loci, we used this to test evidence of replication across sets of loci. As a benchmark, we computed the minimum p-value in 1000 random windows of 50Kb for each study. Less than 5% of random windows had a minimum p<10^−3^ for the non-European ancestry studies.

### Trans-ethnic genetic correlations

We used the popcorn software^30^ to estimate trans-ethnic genetic correlations between studies while accounting for differences in LD structure. This provides an indication of the correlation of causal-variant effect sizes across the genome at SNPs common in both populations. Variant LD scores were estimated for ancestry-matched 1000 Genomes data for each study combination. The estimation of LD scores failed for chromosome 6 for some groups. We therefore left out chromosome 6 from all comparisons. Variants with imputation accuracy r^2^<0.8 or MAF<0.01 were excluded. Popcorn did not converge for any of the studies with less than 20,000 samples. Therefore, results are presented for comparisons between GLGC2013, CKB and BBJ. We estimated effect rather than impact correlations.

### Polygenic scores

We created polygenic scores based on the established lipid loci and assessed their associations with lipid levels in UKHLS, the HELIC cohorts, and APCDR-Uganda, as it was not possible to compute trans-ethnic genetic correlations for these studies. For HELIC and UKHLS, extreme values (*μ* ± 3 *SD*, sex stratified) were filtered. Age, age^2^ and sex were adjusted for by regressing them on the biomarker values and using the residuals as outcomes for subsequent analyses. For each biomarker in each sample set, we checked normality and homoscedasticity. HDL and LDL were approximately normally distributed. For TG levels, a Box Cox transformation was used to normalize the data. APCDR-Uganda phenotype data were rank-based inverse normally transformed.

To make sure PRS were comparable across studies, we excluded variants that were absent, rare (MAF<0.01) or badly imputed (r^2^<0.8) in any of the studies and variants that had different alleles from those in the GLGC. The variant with larger discovery p-value from each correlated pair of SNPs (r^2^>0.1) was also removed. This left 120, 103 and 101 variants for HDL, LDL and TG, respectively. We created trait-specific weighted PRS. The β-regression coefficients from SNP-trait associations in GLGC2017^15^ were used as weights. All biomarkers and scores were scaled to mean=0 and standard deviation=1 for each study so that the regression coefficient represent estimates of the correlation between scores and biomarkers.

We carried out association analyses between each polygenic score and each biomarkers using a linear mixed model with random polygenic effect implemented in GEMMA^38^ in order to account for relatedness and population structure. We used a Bonferroni correction to adjust for multiple testing of three PRS with three different biomarker outcomes (p<0.05/9=0.0056).

### Trans-ethnic colocalization

Differences in allele frequency, LD structure and sample size make it difficult to assess whether a given GWAS hit replicates in samples with different ancestries. Therefore, we applied trans-ethnic colocalization. Colocalization methods test whether the associations in two studies can be explained by the same underlying signal even if the specific causal variant is unknown. The joint likelihood mapping (JLIM) statistic was developed by Chun and colleagues to estimate the posterior probabilities for colocalization between GWAS and eQTL signals and compare them to probabilities of distinct causal variants^16^:

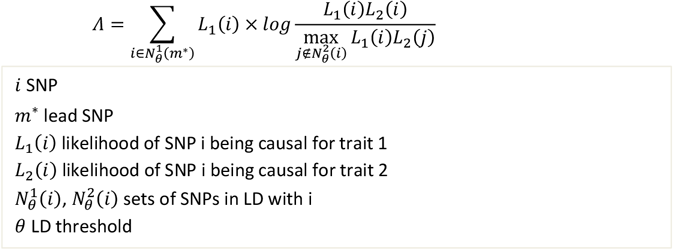

JLIM explicitly accounts for LD structure. Therefore, we assessed whether it is suitable for trans-ethnic colocalization. For samples with summary statistics, LD scores were estimated using ancestry matched samples from the 1000 Genomes Project v3. JLIM assumes only one causal variant within a region in each study. We therefore used a small windows of 50Kb for each known locus to minimise the risk of interference from additional association signals. Distinct causal variants were defined by separation in LD space by r^2^≥0.8 from each other. We excluded loci within the major histocompatibility region due to its complex LD structure. We used a significance threshold of p<0.05 given the evidence of association of the established lipid loci in Europeans and the overall evidence for shared causal genetic architecture across populations for most lipid traits from our other analyses. We compared each target study to UKHLS because of their high level of homogeneity in terms of ancestry, biomarker quantification and study design.

### Simulation

To test the power of trans-ethnic colocalization to detect associations shared between pairs of populations with different ancestry, we ran JLIM on two sets of simulated traits with realistic effect size and environmental noise level. The first set of simulations used the same causal variant in both populations, whereas the second set of simulations discordant causal variants, chosen randomly. We sampled 10,000 randomly chosen biallelic variants with MAF>0.05 and simulated random phenotypes in CKB, APCDR-Uganda und 50,000 individuals with British ancestry from UK Biobank. For each data set relatives were excluded. We also sub-sampled CKB to match the smaller number of individuals in APCDR-Uganda in order to test whether different performance is due to ancestry or sample size. We used a simple linear model to generate the phenotype for each individual i:

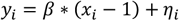

where y is the phenotype value, β is the effect size, x is the number of the alternate alleles carried at the locus and *η_i_*~*N*(0, *σ*^2^), where σ^2^ is the variance of the environmental noise and *Cov*(*η_i_, η_j_*) = 0. We used β= 0.25 and σ^2^= 1.

### Comparison of replicating with non-replicating loci

We aimed to assess whether there are systematic differences between loci that are shared between European ancestry samples and APCDR-Uganda and loci that are not. We identified all loci with evidence of replication based on the above definition that also had significant (p<0.05) colocalization. We only kept one variant per region. We contrasted them with loci where none of the evidence suggested replication: p>0.05 for colocalization, no variant with a lipid association at p<10^−3^ in the region and the lead variant from the discovery study was not rare in APCDR-Uganda. We identified the nearest protein coding gene for each locus and carried out a pathway analyses for the two sets using FUMA^40^. We also assessed the associations of the lead variants with body mass index (BMI) in European ancestry samples using results from a meta-analysis between the GIANT consortium and UK Biobank^17^. We used a Bonferroni adjusted p-value threshold.

### Data availability

The UKHLS EGA accession number is EGAD00010000918. Genotype-phenotype data access for UKHLS is available by application to Metadac (www.metadac.ac.uk). eMERGE is available through dbgap (study ID: phs000888.v1.p1). Summary statistics for GLGC (http://csg.sph.umich.edu/abecasis/public/) and Biobank Japan (http://jenger.riken.jp/en/) are publicly available. The HELIC genotype and WGS datasets have been deposited to the European Genome-phenome Archive (https://www.ebi.ac.uk/ega/home): EGAD00010000518; EGAD00010000522; EGAD00010000610; EGAD00001001636, EGAD00001001637. The APCDR committees are responsible for curation, storage, and sharing of the APCDR-Uganda data under managed access. The array and sequence data have been deposited at the European Genome-phenome Archive (EGA, http://www.ebi.ac.uk/ega/, study accession number EGAS00001000545, datasets EGAS00001001558 and EGAD00001001639 respectively) and can be requested through. Requests for access to phenotype data may be directed to.

## Supplemental data

The supplemental data contain three figures and two tables.

## Funding and acknowledgements

We would like to thank the participants and volunteers of the each of the studies. We appreciate the advice received from Brielin Brown regarding the use of Popcorn and from Shamil Sunyaev and Sung Chun regarding the suitability of JLIM for trans-ethnic colocalization.

This work was funded by the Wellcome Trust (WT098051), (212360/Z/18/Z), and the European Research Council (ERC-2011-StG 280559-SEPI). The MANOLIS study is dedicated to the memory of Manolis Giannakakis, 1978-2010. We thank the residents of the Pomak villages and of the Mylopotamos villages for taking part. The HELIC study has been supported by many individuals who have contributed to sample collection (including Antonis Athanasiadis, Olina Balafouti, Christina Batzaki, Georgios Daskalakis, Eleni Emmanouil, Pounar Feritoglou, Chrisoula Giannakaki, Margarita Giannakopoulou, Kiki Kaldaridou, Anastasia Kaparou, Vasiliki Kariakli, Stella Koinaki, Dimitra Kokori, Maria Konidari, Hara Koundouraki, Dimitris Koutoukidis, Vasiliki Mamakou, Eirini Mamalaki, Eirini Mpamiaki, Nilden Selim, Nesse Souloglou, Maria Tsoukana, Dimitra Tzakou, Katerina Vosdogianni, Niovi Xenaki, Eleni Zengini), data entry (Thanos Antonos, Dimitra Papagrigoriou, Betty Spiliopoulou), sample logistics (Sarah Edkins, Emma Gray), genotyping (Suzannah Bumpstead, Robert Andrews, Hannah Blackburn, Doug Simpkin, Siobhan Whitehead), research administration (Anja Kolb-Kokocinski, Carol Smee, Danielle Walker) and informatics (Kathleen Stirrups, Martin Pollard, Josh Randall).

APCDR-Uganda was funded by the Wellcome Trust, The Wellcome Trust Sanger Institute (WT098051), the UK Medical Research Council (G0901213-92157, G0801566, and MR/K013491/1), and the Medical Research Council/Uganda Virus Research Institute Uganda Research Unit on AIDS core funding. We thank the African Partnership for Chronic Disease Research (APCDR) for providing a network to support this study as well as a repository for deposition of curated data. We also thank all study participants who contributed to this study.

The UK Household Longitudinal Study was funded by grants from the Economic & Social Research Council (ES/H029745/1) and the Wellcome Trust (WT098051). UKHLS is led by the Institute for Social and Economic Research at the University of Essex and funded by the Economic and Social Research Council. The survey was conducted by NatCen and the genome-wide scan data were analysed and deposited by the Wellcome Trust Sanger Institute.

The Understanding Society Scientific Group are Michaela Benzeval(1), Jonathan Burton(1), Nicholas Buck(1), Annette Jäckle(1), Meena Kumari(1), Heather Laurie(1), Peter Lynn(1), Stephen Pudney(1), Birgitta Rabe(1), Dieter Wolke(2)

1. Institute for Social and Economic Research
2. University of Warwick

## Web resources

We have made code to run trans-ethnic colocalization using JLIM and simulations available through github: https://github.com/KarolineKuchenbaecker/TEColoc

