## Supplementary Tables and Figures for "The transferability of lipid loci across African, Asian and European cohorts"

### Supplementary figures and tables

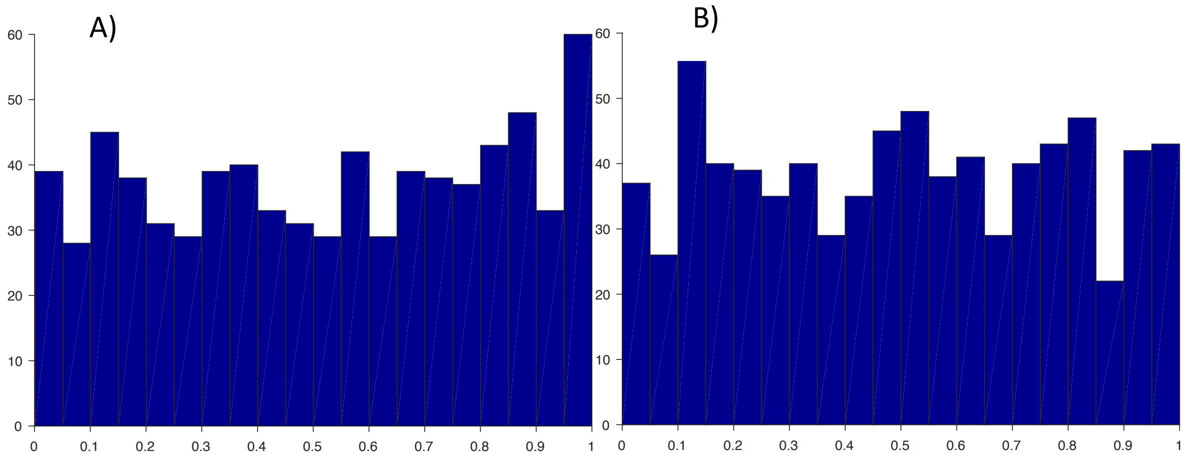


**Supplementary Figure 1:** Distribution of p-values from the trans-ethnic colocalization for simulated traits assuming distinct causal variants in UK Biobank and A) CKB and B) APCDR-Uganda

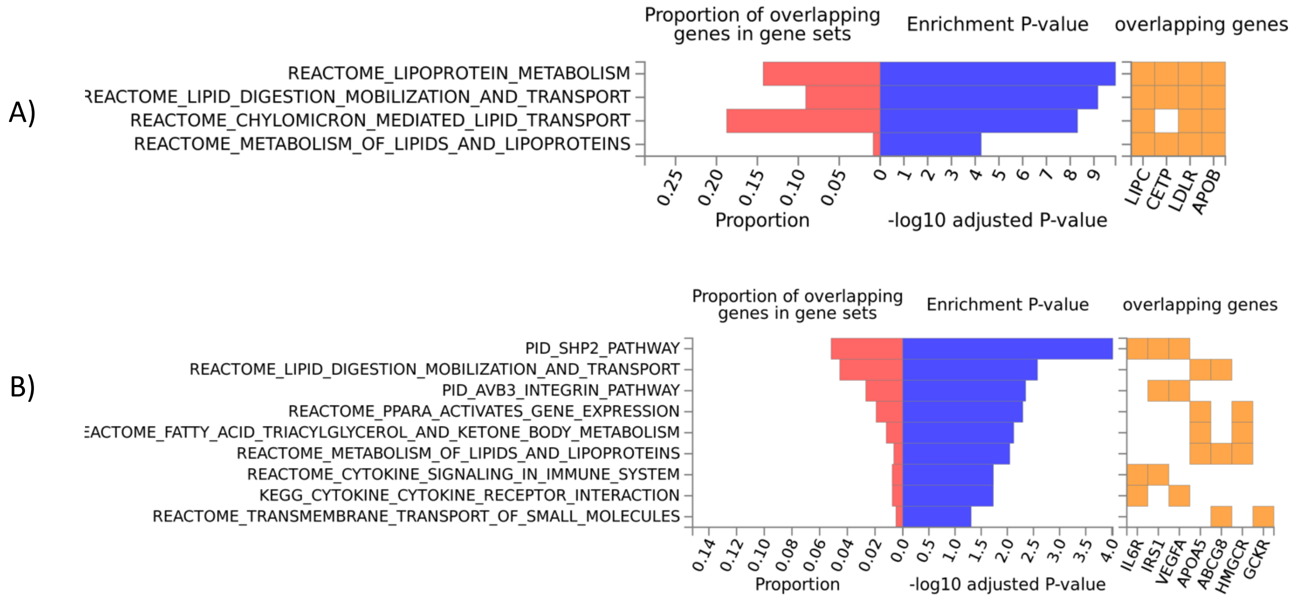


**Supplementary Figure 2: Enrichment of canonical pathways** (MsigDB c2) for genes proximal to established lipid-associated SNPs that A) replicate in APCDR-Uganda and B) do not replicate in APCDR-Uganda

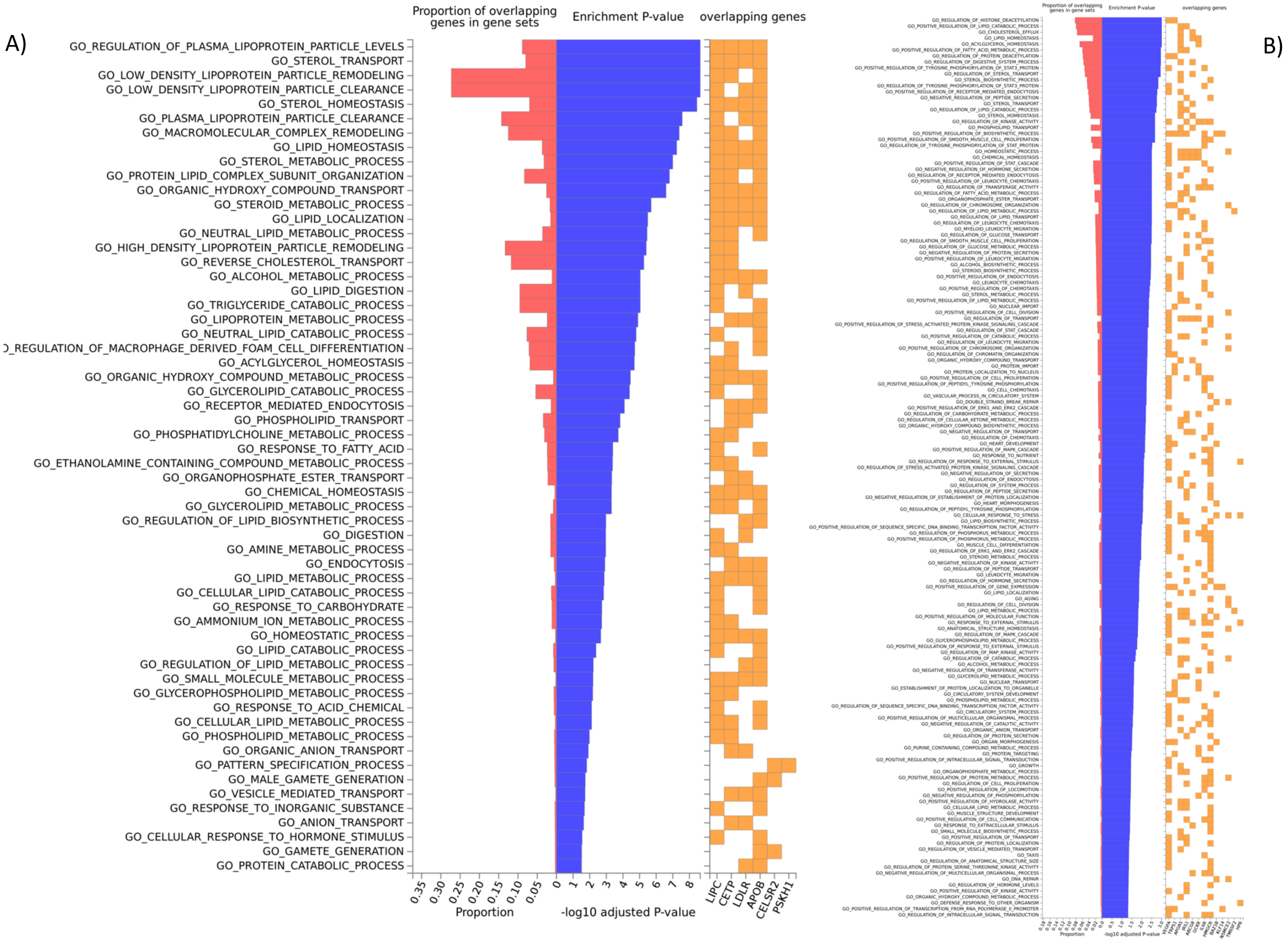
A)
Proportion of overlapping genes in gene sets
Enrichment P-value
overlapping genes
GO_REGULATION_OF_PLASMA_LIPOPROTEIN_PARTICLE_LEVELS
GO_STEROL_TRANSPORT
GO_LOW_DENSITY_LIPOPROTEIN_PARTICLE_REMODELING
GO_LOW_DENSITY_LIPOPROTEIN_PARTICLE_CLEARANCE
GO_STEROL_HOMEOSTASIS
GO_PLASMA_LIPOPROTEIN_PARTICLE_CLEARANCE
GO_MACROMOLECULAR_COMPLEX_REMODELING
GO_LIPID_HOMEOSTASIS
GO_STEROL_METABOLIC_PROCESS
GO_PROTEIN_LIPID_COMPLEX_SUBUNIT_ORGANIZATION
GO_ORGANIC_HYDROXY_COMPOUND_TRANSPORT
GO_STEROID_METABOLIC_PROCESS
GO_LIPID_LOCALIZATION
GO_NEUTRAL_LIPID_METABOLIC_PROCESS
GO_HIGH_DENSITY_LIPOPROTEIN_PARTICLE_REMODELING
GO_REVERSE_CHOLESTEROL_TRANSPORT
GO_ALCOHOL_METABOLIC_PROCESS
GO_LIPID_DIGESTION
GO_TRIGLYCERIDE_CATABOLIC_PROCESS
GO_LIPOPROTEIN_METABOLIC_PROCESS
GO_NEUTRAL_LIPID_CATABOLIC_PROCESS
_REGULATION_OF_MACROPHAGE_DERIVED_FOAM_CELL_DIFFERENTIATION
GO_ACYLGLYCEROL_HOMEOSTASIS
GO_ORGANIC_HYDROXY_COMPOUND_METABOLIC_PROCESS
GO_GLYCEROLIPID_CATABOLIC_PROCESS
GO_RECEPTOR_MEDIATED_ENDOCYTOSIS
GO_PHOSPHOLIPID_TRANSPORT
GO_PHOSPHATIDYLCHOLINE_METABOLIC_PROCESS
GO_RESPONSE_TO_FATTY_ACID
GO_ETHANOLAMINE_CONTAINING_COMPOUND_METABOLIC_PROCESS
GO_ORGANOPHOSPHATE_ESTER_TRANSPORT
GO_CHEMICAL_HOMEOSTASIS
GO_GLYCEROLIPID_METABOLIC_PROCESS
GO_REGULATION_OF_LIPID_BIOSYNTHETIC_PROCESS
GO_DIGESTION
GO_AMINE_METABOLIC_PROCESS
GO_ENDOCYTOSIS
GO_LIPID_METABOLIC_PROCESS
GO_CELLULAR_LIPID_CATABOLIC_PROCESS
GO_RESPONSE_TO_CARBOHYDRATE
GO_AMMONIUM_ION_METABOLIC_PROCESS
GO_HOMEOSTATIC_PROCESS
GO_LIPID_CATABOLIC_PROCESS
GO_REGULATION_OF_LIPID_METABOLIC_PROCESS
GO_SMALL_MOLECULE_METABOLIC_PROCESS
GO_GLYCEROPHOSPHOLIPID_METABOLIC_PROCESS
GO_RESPONSE_TO_ACID_CHEMICAL
GO_CELLULAR_LIPID_METABOLIC_PROCESS
GO_PHOSPHOLIPID_METABOLIC_PROCESS
GO_ORGANIC_ANION_TRANSPORT
GO_PATTERN_SPECIFICATION_PROCESS
GO_MALE_GAMETE_GENERATION
GO_VESICLE_MEDIATED_TRANSPORT
GO_RESPONSE_TO_INORGANIC_SUBSTANCE
GO_ANION_TRANSPORT
GO_CELLULAR_RESPONSE_TO_HORMONE_STIMULUS
GO_GAMETE_GENERATION
GO_PROTEIN_CATABOLIC_PROCESS
0.35 0.30 0.25 0.20 0.15 0.10 0.05 0
Proportion
0 1 2 3 4 5 6 7 8
-log10 adjusted P-value
LIPC
CETP
LDLR
APOB
CELSR2
PSKH1
B)

**Supplementary Figure 3: Enrichment of GO biological processes** (MsigDB c5) for genes proximal to established lipid-associated SNPs that A) replicate in APCDR-Uganda and B) do not replicate in APCDR-Uganda

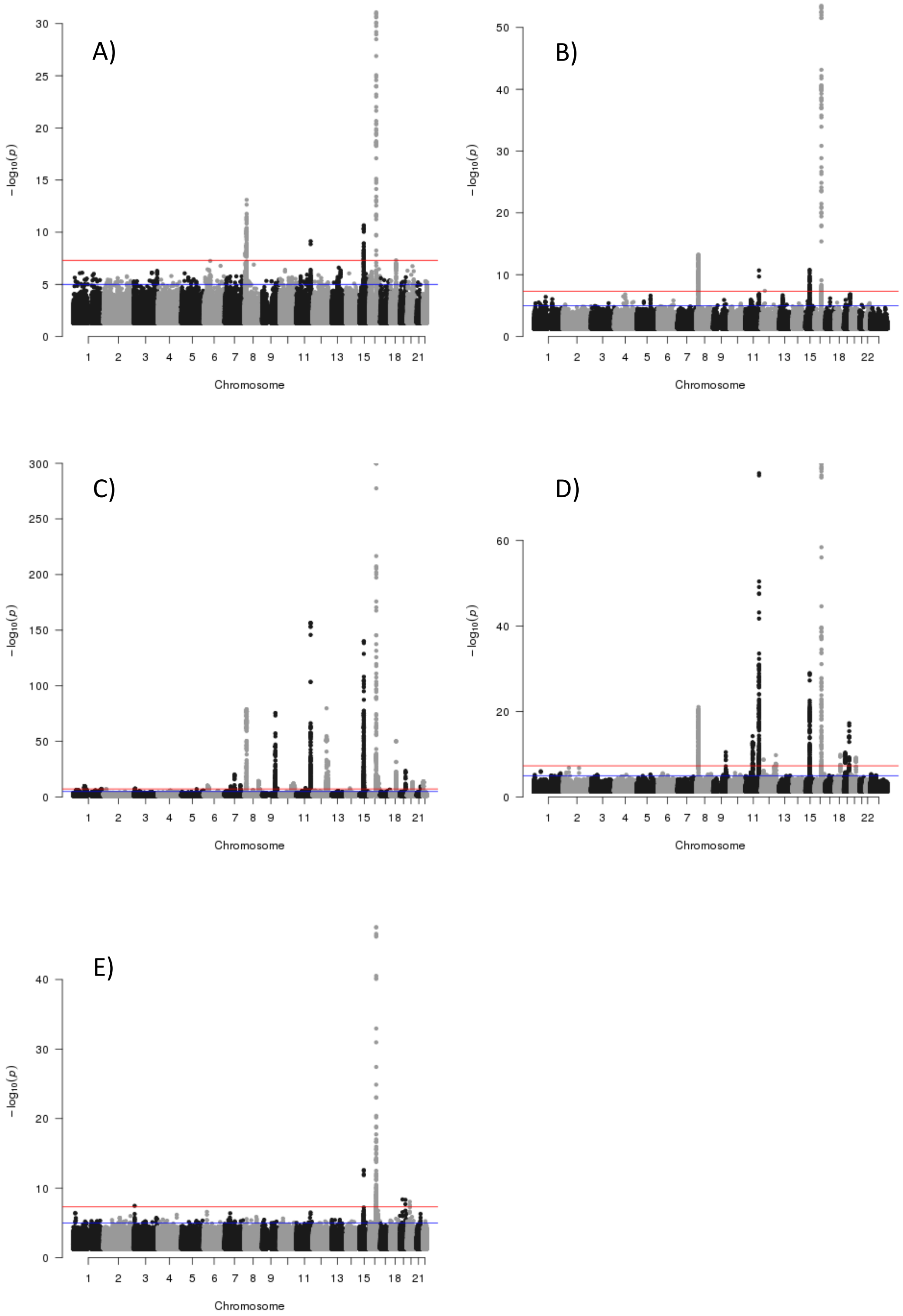


**Supplementary Figure 4: Manhattan plot** for SNP associations with HDL cholesterol levels, for A) eMerge, B) UKHLS, C) Biobank Japan, D) China Kadoorie Biobank, E) APCDR-Uganda

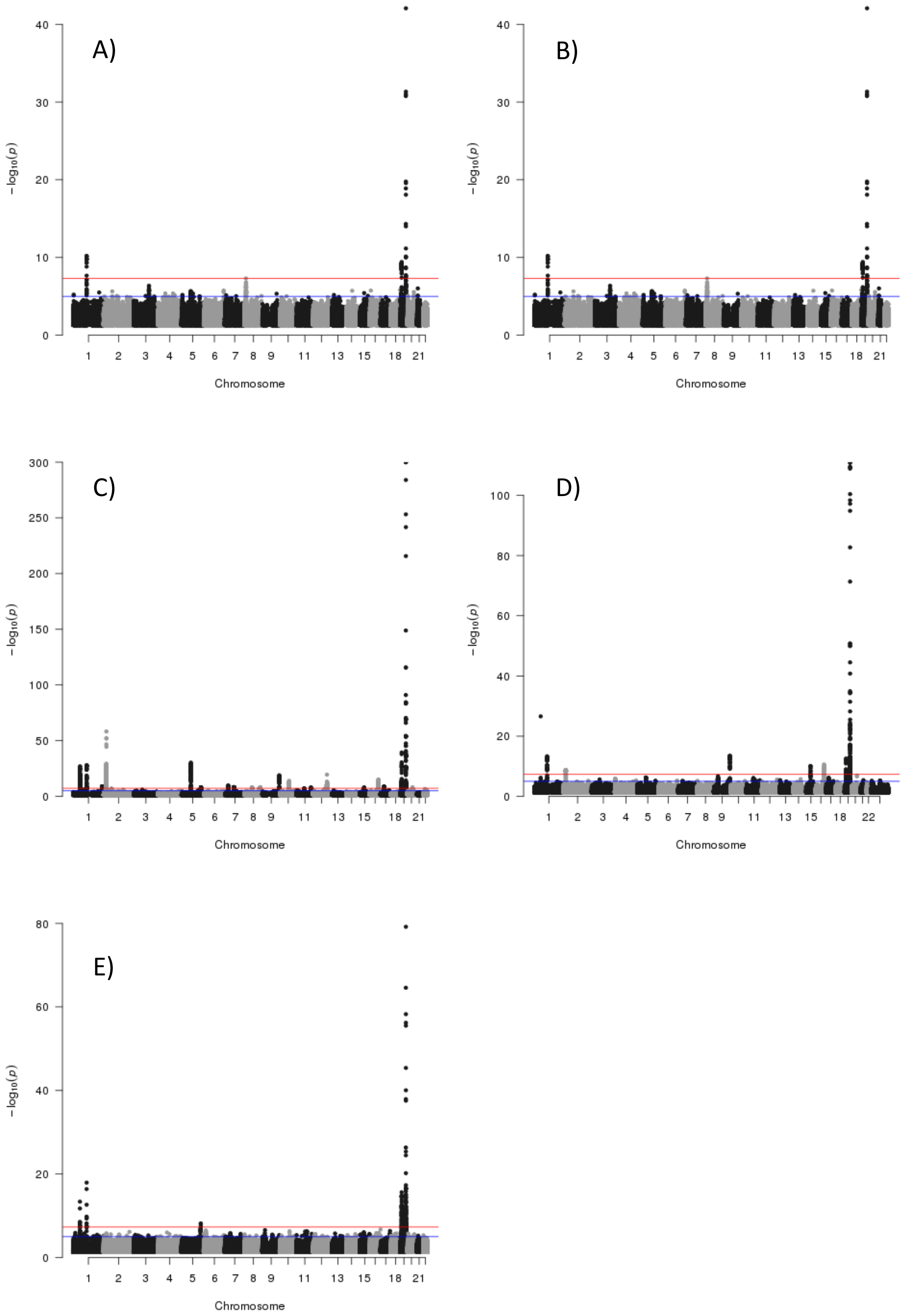


**Supplementary Figure 5: Manhattan plot** for SNP associations with LDL cholesterol levels, for A) eMerge, B) UKHLS, C) Biobank Japan, D) China Kadoorie Biobank, E) APCDR-Uganda

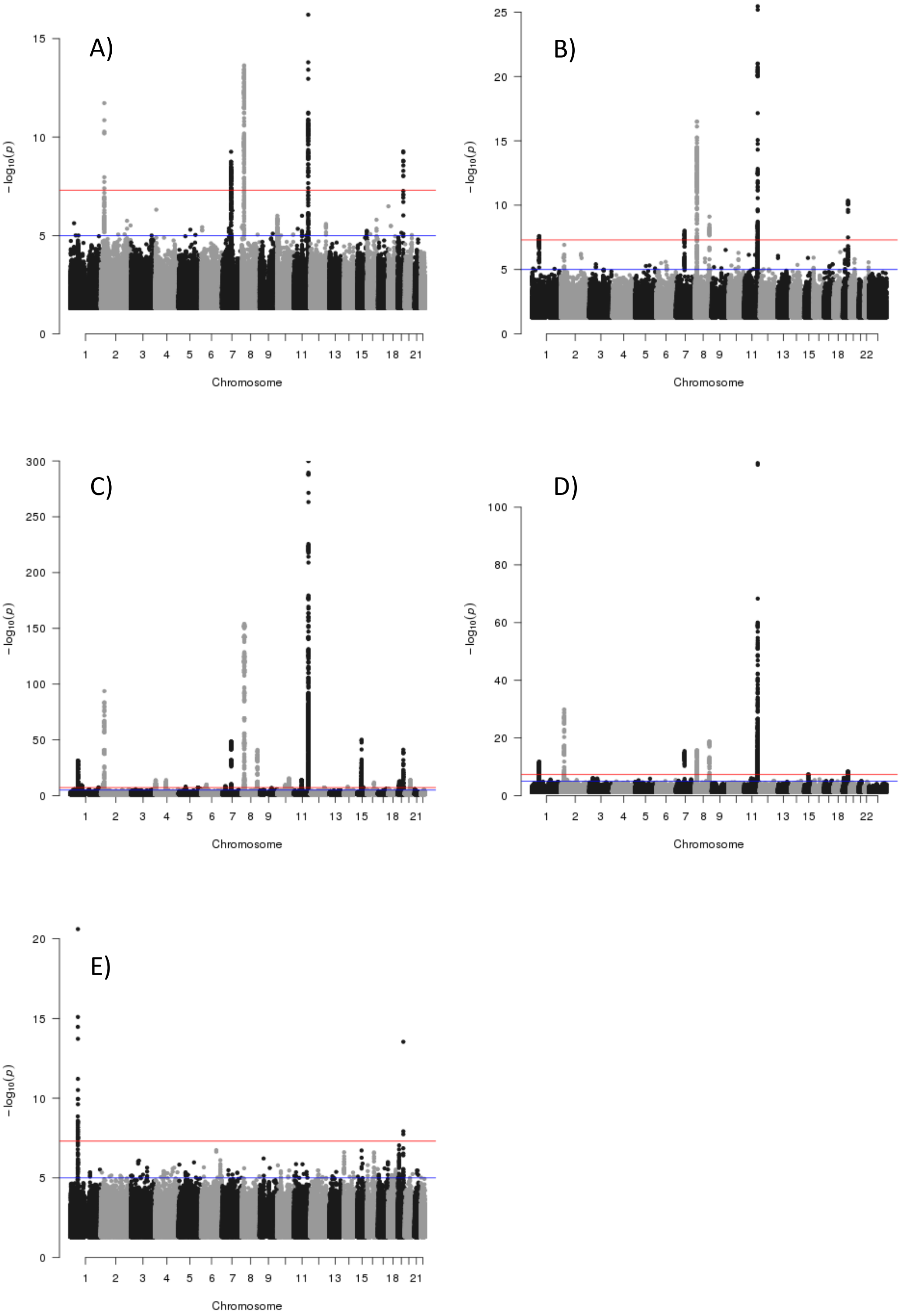


**Supplementary Figure 6: Manhattan plot** for SNP associations with triglyceride levels, for A) eMerge, B) UKHLS, C) Biobank Japan, D) China Kadoorie Biobank, E) APCDR-Uganda

**Supplementary Table 1: Trans-ethnic genetic correlation** estimates and p-values for a test whether the genetic correlation is 1 for each lipid biomarker in one study with each lipid biomarkers in the other study

| biomarker | correlation | standard error | p-value* |
|---|---|---|---|
| **GLGC2013 (European) – China Kadoorie Biobank** | | | |
| HDL-HDL | 0.999 | - | - |
| LDL-LDL | 0.778 | 0.300 | 0.460 |
| TG-TG | 0.999 | - | - |
| HDL - LDL | 0.068 | 0.229 | 0.000 |
| HDL - TG | -0.550 | 0.176 | 0.000 |
| LDL - HDL | -0.238 | 0.143 | 0.000 |
| LDL - TG | 0.473 | 0.155 | 0.001 |
| TG - HDL | -0.741 | 0.162 | 0.000 |
| TG - LDL | -0.226 | 0.195 | 0.000 |
| **GLGC2013 (European) – Biobank Japan** | | | |
| HDL-HDL | 0.999 | 0.081 | 0.999 |
| LDL-LDL | 0.959 | 0.138 | 0.765 |
| TG-TG | 0.961 | 0.066 | 0.555 |
| HDL - LDL | -0.055 | 0.097 | 0.000 |
| HDL - TG | -0.592 | 0.130 | 0.000 |
| LDL - HDL | -0.277 | 0.150 | 0.000 |
| LDL - TG | 0.294 | 0.056 | 0.000 |
| TG - HDL | -0.481 | 0.176 | 0.000 |
| TG - LDL | -0.038 | 0.091 | 0.000 |
| **China Kadoorie Biobank – Biobank Japan** | | | |
| HDL-HDL | 0.999 | - | - |
| LDL-LDL | 0.871 | 0.225 | 0.566 |
| TG-TG | 0.999 | - | - |
| HDL - LDL | 0.085 | 0.141 | 0.000 |
| HDL - TG | -0.618 | 0.151 | 0.000 |
| LDL - HDL | 0.290 | 0.185 | 0.000 |
| LDL - TG | 0.180 | 0.169 | 0.000 |
| TG - HDL | -0.862 | 0.186 | 0.000 |
| TG - LDL | -0.097 | 0.153 | 0.000 |

* When the estimate is close to the boundary of 1, popcorn cannot compute the standard error and p-value

**Supplementary Table 2: Associations of polygenic scores** based on established lipid-associated loci and levels of each of the other lipid biomarkers in UKHLS, HELIC-MANOLIS, -Pomak, and APCDR-Uganda using a linear mixed model analysis.

| score - trait | p-value | correlation | SE (β) |
|---|---|---|---|
| **APCDR-Uganda** | | | |
| HDL - LDL | 2.34E-01 | -1.49E-02 | 1.25E-02 |
| HDL - TG | 2.49E-01 | -1.44E-02 | 1.25E-02 |
| LDL - HDL | 6.75E-10 | -7.73E-02 | 1.25E-02 |
| LDL - TG | 7.27E-04 | 4.22E-02 | 1.25E-02 |
| TG - HDL | 6.81E-01 | -5.13E-03 | 1.25E-02 |
| TG - LDL | 1.86E-04 | -4.67E-02 | 1.25E-02 |
| **UKHLS** | | | |
| HDL - LDL | 2.02E-07 | -5.26E-02 | 1.01E-02 |
| HDL - TG | 1.63E-12 | -7.20E-02 | 1.02E-02 |
| LDL - HDL | 3.97E-07 | -5.16E-02 | 1.01E-02 |
| LDL - TG | 4.07E-01 | 8.45E-03 | 1.02E-02 |
| TG - HDL | 1.67E-39 | -1.34E-01 | 1.01E-02 |
| TG - LDL | 2.47E-01 | 1.17E-02 | 1.01E-02 |
| **HELIC-Pomak** | | | |
| HDL - LDL | 1.97E-02 | -7.22E-02 | 3.09E-02 |
| HDL - TG | 4.96E-05 | -1.28E-01 | 3.10E-02 |
| LDL - HDL | 6.00E-02 | -6.08E-02 | 3.18E-02 |
| LDL - TG | 9.18E-01 | 3.43E-03 | 3.19E-02 |
| TG - HDL | 6.16E-07 | -1.57E-01 | 3.05E-02 |
| TG - LDL | 1.70E-02 | 7.39E-02 | 3.08E-02 |
| **HELIC-MANOLIS** | | | |
| HDL - LDL | 7.56E-01 | 9.19E-03 | 2.96E-02 |
| HDL - TG | 2.13E-03 | -9.36E-02 | 3.01E-02 |
| LDL - HDL | 4.42E-01 | -2.29E-02 | 2.97E-02 |
| LDL - TG | 9.64E-01 | -1.36E-03 | 3.03E-02 |
| TG - HDL | 1.25E-05 | -1.33E-01 | 2.98E-02 |
| TG - LDL | 9.56E-01 | -1.63E-03 | 2.98E-02 |

**Supplementary Table 3. P-value for the trans-ethnic colocalization** based on the JLIM model for established lipid-associated loci in UKHLS, China Kadoorie Biobank (CKB), Biobank Japan (BBJ) and Uganda (APCDR).

| | | | | | GLGC | | replication[a] | | | | JLIM p-value[b] | | |
|---|---|---|---|---|---|---|---|---|---|---|---|---|---|
| rs-id | chr | position | near gene | MAF | p-value | Multi[c] | UKHLS | CKB | BBJ | APCDR | CKB | BBJ | APCDR |
| **HDL** | | | | | | | | | | | | | |
| rs4660293 | 1 | 40028180 | PABPC4 | 0.21 | 6.1E-36 | distant | uc | ns | cor | ns | 0.8 | 0.014 | NA |
| rs11755393 | 6 | 34824636 | UHRF1BP1 | 0.36 | 4.2E-23 | distant | ns | ns | cor | ns | 0.035 | 0 | NA |
| rs1178979 | 7 | 72856430 | BAZ1B | 0.18 | 1.3E-26 | near | uc | uc | cor | ns | 0 | 0 | NA |
| rs4731702 | 7 | 130433384 | KLF14 | 0.46 | 1.2E-35 | distant | ns | cor | cor | ns | 0.14 | 0.003 | NA |
| rs4841132 | 8 | 9183596 | PPP1R3B | 0.9 | 1.0E-123 | no | ns | ns | ns | cor | 0.99 | 0.24 | 0.16 |
| rs328 | 8 | 19819724 | LPL | 0.098 | 1.7E-316 | near | cor | cor | cor | cor | 0.21 | 0.005 | 0.62 |
| rs2954033 | 8 | 126493746 | NSMCE2 | 0.72 | 3.0E-61 | near | cor | cor | cor | ns | 0.94 | 0.002 | NA |
| rs643531 | 9 | 15296034 | TTC39B | 0.88 | 3.8E-42 | no | ns | ns | uc | uc | NA | NA | NA |
| rs2066714 | 9 | 107586753 | ABCA1 | 0.15 | 3.6E-31 | near | uc | cor | cor | uc | 0.97 | 0.007 | 0.94 |
| rs1883025 | 9 | 107664301 | ABCA1 | 0.26 | 2.1E-118 | near | uc | cor | cor | uc | 0.84 | 0.8 | 0.97 |
| rs2792751 | 10 | 113940329 | GPAM | 0.73 | 3.8E-21 | near | ns | ns | cor | uc | 0.002 | 0.002 | NA |
| rs7350481 | 11 | 116586283 | APOA5 | 0.91 | 3.2E-100 | distant | cor | cor | cor | uc | 0 | 0 | 0.99 |
| rs964184 | 11 | 116648917 | ZPR1 | 0.85 | 2.6E-217 | near | cor | cor | cor | uc | 1 | 1 | 1 |
| rs10468017 | 15 | 58678512 | LIPC | 0.27 | 1.8E-306 | near | cor | cor | cor | cor | 1 | 0.98 | 0.99 |
| rs1800588 | 15 | 58723675 | LIPC | 0.24 | 0 | distant | cor | cor | cor | cor | 0.007 | 0.009 | 0.017 |
| rs247616 | 16 | 56989590 | CETP | 0.31 | 0 | near | cor | cor | cor | cor | 0 | 0 | 0 |
| rs3764261 | 16 | 56993324 | CETP | 0.31 | 0 | near | cor | cor | cor | cor | 0 | 0 | 0 |
| rs34065661 | 16 | 56995935 | CETP | 0.005 | 5.6E-103 | near | uc | uc | uc | cor | 0 | 0 | 0 |
| rs16942887 | 16 | 67928042 | PSKH1 | 0.13 | 9.8E-93 | near | ns | ns | cor | cor | 0.96 | 0.29 | 0.025 |
| rs72836561 | 17 | 41926126 | CD300LG | 0.028 | 8.1E-111 | no | uc | uc | uc | ns | NA | NA | NA |
| rs7241918 | 18 | 47160953 | LIPG | 0.85 | 1.2E-104 | distant | cor | cor | cor | ns | 0.16 | 1 | 1 |
| rs116843064 | 19 | 8429323 | ANGPTL4 | 0.02 | 4.8E-146 | near | uc | ns | ns | uc | NA | NA | NA |
| rs769449 | 19 | 45410002 | APOE | 0.11 | 6.9E-129 | near | cor | cor | cor | cor | 0.009 | 0.02 | 0.95 |
| rs386000 | 19 | 54792761 | LILRB2 | 0.22 | 1.1E-41 | distant | cor | uc | cor | cor | 0 | 1 | 0.71 |
| **LDL** | | | | | | | | | | | | | |
| rs11591147 | 1 | 55505647 | PCSK9 | 0.015 | 0.0 | near | uc | uc | uc | uc | 0.94 | 0.64 | 0.92 |
| rs12740374 | 1 | 109817590 | CELSR2 | 0.22 | 0.0 | near | cor | cor | cor | cor | 0 | 0 | 0 |

| | | | | | | | | | | | | | |
|---|---|---|---|---|---|---|---|---|---|---|---|---|---|
| rs1367117 | 2 | 21263900 | APOB | 0.28 | 3.6E-278 | near | cor | cor | cor | cor | 1 | 1 | 0 |
| rs541041 | 2 | 21294975 | APOB | 0.81 | 1.3E-287 | distant | cor | uc | uc | cor | 1 | 1 | NA |
| rs4245791 | 2 | 44074431 | ABCG8 | 0.72 | 1.7E-120 | near | uc | ns | ns | ns | NA | 0.81 | 0.99 |
| rs3846662 | 5 | 74651084 | HMGCR | 0.48 | 3.3E-128 | near | cor | cor | cor | ns | 0.002 | 0 | 1 |
| rs2737229 | 8 | 116648565 | TRPS1 | 0.34 | 8.9E-15 | distant | cor | ns | cor | ns | 0.015 | 0.025 | NA |
| rs635634 | 9 | 136155000 | IL6R | 0.19 | 4.9E-109 | near | ns | cor | cor | ns | 0.96 | 0.97 | NA |
| rs2000999 | 16 | 72108093 | HPR | 0.2 | 4.0E-71 | distant | cor | cor | cor | ns | 1 | 0 | 1 |
| rs6511720 | 19 | 11202306 | LDLR | 0.11 | 0.0 | near | cor | uc | uc | cor | 0 | NA | 0 |
| rs28399654 | 19 | 45316588 | BCAM | 0.027 | 7.5E-232 | distant | cor | uc | uc | cor | 1 | 1 | NA |
| rs7412 | 19 | 45412079 | APOE | 0.075 | 0.0E+00 | near | cor | cor | cor | cor | 0 | 0 | 0 |
| **Triglycerides** | | | | | | | | | | | | | |
| rs10889353 | 1 | 63118196 | DOCK7 | 0.33 | 6.4E-170 | no | cor | cor | cor | uc | 0 | 0 | 0.88 |
| rs676210 | 2 | 21231524 | APOB | 0.26 | 4.9E-118 | near | cor | uc | cor | uc | 1 | 1 | 1 |
| rs1260326 | 2 | 27730940 | GCKR | 0.63 | 0.0 | near | cor | cor | cor | ns | 0 | NA | NA |
| rs2943641 | 2 | 227093745 | IRS1 | 0.66 | 4.9E-33 | no | ns | ns | cor | ns | 0.006 | NA | 1 |
| rs6905288 | 6 | 43758873 | VEGFA | 0.59 | 9.0E-35 | near | cor | ns | cor | ns | 0 | 0 | NA |
| rs1178979 | 7 | 72856430 | BAZ1B | 0.18 | 1.5E-179 | near | cor | cor | cor | ns | 0 | 0 | NA |
| rs35332062 | 7 | 73012042 | MLXIPL | 0.12 | 5.2E-205 | distant | cor | cor | cor | uc | 0.99 | 1 | NA |
| rs326 | 8 | 19819439 | LPL | 0.3 | 0.0 | near | cor | cor | cor | uc | 0.91 | 0.91 | 0.72 |
| rs2954029 | 8 | 126490972 | TRIB1 | 0.45 | 8.3E-205 | near | cor | cor | cor | ns | 0 | 0 | 1 |
| rs1883025 | 9 | 107664301 | ABCA1 | 0.26 | 1.2E-13 | no | uc | ns | ns | ns | 1 | 0.001 | NA |
| rs7350481 | 11 | 116586283 | APOA5 | 0.91 | 0.0 | distant | cor | cor | cor | uc | 0 | 0 | 0 |
| rs11820589 | 11 | 116633862 | APOA5 | 0.066 | 4.4E-133 | near | cor | uc | uc | ns | 0 | 1 | 1 |
| rs2075291 | 11 | 116661392 | APOA5 | 0.003 | 5.7E-65 | near | uc | cor | cor | ns | NA | 1 | NA |
| rs10047462 | 11 | 116722041 | SIK3 | 0.86 | 9.9E-180 | near | cor | cor | cor | uc | 0 | 0 | 1 |
| rs247616 | 16 | 56989590 | CETP | 0.31 | 2.4E-38 | near | ns | ns | cor | cor | 0.014 | 0.024 | 0.78 |
| rs116843064 | 19 | 8429323 | ANGPTL4 | 0.02 | 4.2E-175 | near | uc | ns | ns | ns | NA | NA | NA |
| rs58542926 | 19 | 19379549 | TM6SF2 | 0.074 | 3.7E-125 | no | cor | cor | cor | ns | NA | NA | NA |
| rs439401 | 19 | 45414451 | APOE | 0.63 | 2.7E-168 | near | cor | cor | cor | uc | 0.009 | 0.53 | 1 |

[a] indicates whether any variant from the credible set ("cor") or any uncorrelated variant within 50kb ("uc") is associated with the target biomarker at $p<10^{-3}$ in each of the target studies

[b] p-value from the JLIM trans-ethnic colocalization analysis using UKHLS as the comparison set

[c] indicates whether multiple independent hits have been reported within 50kb ("near") or 1Mb ("distant")

**Supplementary Table 4: Study description** and mean levels of HDL-cholesterol, LDL-cholesterol and triglycerides (TG) in mmol/l

| study | acronym | population | array | N SNPs genotyped, imputed | N samples | %female | mean age | mean HDL | mean LDL | mean TG |
|---|---|---|---|---|---|---|---|---|---|---|
| UK Household Longitudinal Study[1] | **UKHLS** | British | HumanCoreExome | 248K, 26M | 9,962 | 44 | 52 | 1.55 | 3.02 | 1.81 |
| electronic MEdical Records and GEnomics[2] | **eMERGE** | European ancestry | GWAS arrays, Metabochip | 17.6M | 14,528 | 43 | 61 | 1.33 | 2.80 | 1.77 |
| Global Lipids Genetics Consortium[3] | **GLGC2013** (meta-analysis) | European ancestry | 23 studies GWAS arrays, 37 Metabochip | 200K, 2.5M | 188,577 | | | | | |
| Global Lipids Genetics Consortium[4] | **GLGC2017** (meta-analysis) | European ancestry | HumanExome | 242,289 | 237,050 | | | | | |
| African Partnership for Chronic Disease Research - Uganda[5] | **APCDR-Uganda** | Ugandan | HumanOmni2.5 | 2.2M, 20M | 6,407 | 56 | 34 | 1.02 | 2.05 | 1.18 |
| China Kadoorie Biobank[6] | **CKB** | Chinese | Custom Affymetrix Axiom Array | 701K/830K, 10M | 21,295 | 62 | 60 | 1.37 | 2.19 | 1.69 |
| RIKEN Biobank Japan[7] | **BBJ** | Japanese | HumanOmniExpress | 6M | 162,255 | 63 | 43 | 1.42 | 3.38 | 1.50 |
| Hellenic Isolated Cohorts[8,9] | **HELIC-MANOLIS, -Pomak** | Isolated Greek populations | Whole-genome sequencing | 24M | 1,641, 1,945 | 42,34 | 62,45 | 1.28, 1.18 | 3.27, 3.09 | 1.61, 1.58 |

**Study references**
